# Who’s who in the western Hermann’s tortoise conservation: a STR toolkit and reference database for wildlife forensic genetic analyses

**DOI:** 10.1101/484030

**Authors:** Biello Roberto, Zampiglia Mauro, Corti Claudia, Deli Gianluca, Biaggini Marta, Delaugerre Michel, Di Tizio Luciano, Leonetti Francesco Luigi, Olivieri Oliviero, Pellegrino Francesco, Romano Antonio, Sperone Emilio, Trabalza-Marinucci Massimo, Bertorelle Giorgio, Canestrelli Daniele

**Affiliations:** Dipartimento di Scienze della Vita e Biotecnologie, Università di Ferrara, Via Luigi Borsari 46, 44121 Ferrara, Italy; Dipartimento di Scienze Ecologiche e Biologiche, Università della Tuscia, Largo dell’Università s.n.c., 01100 Viterbo, Italy; Museo di Storia Naturale dell’Università di Firenze, Sezione di Zoologia “La Specola”, Via Romana 17, 50125 Firenze, Italy; Dipartimento di Medicina Veterinaria, Università di Perugia, Via San Costanzo 4, 06126 Perugia, Italy; Conservatoire du littoral, Résidence St Marc, 2, rue Juge Falcone, 20200 Bastia, France; Societas Herpetologica Italica, Sezione Abruzzo-Molise, Via Federico Salomone 112, 66100 Chieti, Italy; DiBEST, Università della Calabria, via P. Bucci, 87036, Rende (CS), Italy; MUSE: Museo delle Scienze, Sezione di Zoologia dei Vertebrati, corso del Lavoro e della Scienza 3, 38122 Trento, Italy; CNR-ISAFOM: Consiglio Nazionale delle Ricerche, Istituto per i sistemi agricoli e forestali del Mediterraneo, Via Patacca 85, 80056 Ercolano (NA), Italy

**Keywords:** Wildlife forensic genetics, Pet trade, Illegal animal translocation, Assignment tests, STR toolkit, Mediterranean tortoises, *Testudo hermanni*

## Abstract

Illegal trade is threatening tortoise populations worldwide since decades. Nowadays, however, DNA typing and forensic genetic approaches allow to investigate geographic origin of confiscated animals and to relocate them into the wild, provided that suitable molecular tools and reference data are available. Here we assess the suitability of a small panel of microsatellite markers to investigate patterns of illegal translocations and to assist forensic genetic applications in the endangered Mediterranean land tortoise *Testudo hermanni hermanni*. We used the microsatellite panel to (i) increase the understanding of the population genetic structure in wild populations with new data from previously unsampled geographic areas (overall 461 wild individuals from 28 sampling sites); (ii) detect the presence of non-native individuals in wild populations; and (iii) identify the most likely geographic area of origin of 458 confiscated individuals hosted in Italian seizure and recovery centers. Our analysis initially identified six major genetic clusters corresponding to different geographic macro-areas along the Mediterranean range. Long-distance migrants among wild populations, due to translocations, were found and removed from the reference database. Assignment tests allowed us to allocate approximately 70% of confiscated individuals of unknown origin to one of the six Mediterranean macro-areas. Most of the assigned tortoises belonged to the genetic cluster corresponding to the area where the respective captivity center was located. However, we also found evidence of long-distance origin of confiscated individuals, especially in centers along the Adriatic coast and facing the Balkan regions, a well-known source of illegally traded individuals. Our results clearly show the role for reintroduction projects of the microsatellite panel, which was useful to re-assign most of the confiscated individuals to the respective macro-area of origin. At the same time, the microsatellite panel can assist future forensic genetic applications to detect illegal trade and possess of *Testudo hermanni* individuals.

## INTRODUCTION

Over-collection and illegal trade of wildlife species for consumption or pet market are among the main threats to biodiversity [1], and reptiles currently represent the second most affected vertebrate class, after birds [2,3]. According to [4], the European Union (EU) is the top global importer of live reptiles for the pet trade (valued at €7 million in 2005). Because of this practice, a significant number of reptile populations have already been severely decimated (e.g., [5–8]). Intentional harvest is considered the second largest threat to the survival of many reptile species [9] and, as a consequence, reptiles’ pet trade is strongly restrained by CITES.

Relocating confiscated individuals implies the identification of their natural source areas, which has long been a challenging task in the absence of clear morphological differences among natural populations and the consequent lack of simple diagnostic traits [1]. However, DNA typing and forensic genetic tools are providing straightforward and increasingly appreciated approaches for this purpose, allowing also the identification of hybrids. Noteworthy, the use of these wildlife forensic genetic tools implies the gathering of multiple population genetics information in a single analytic framework, such as the assessment of the genetic variation and its deep population structure at the geographical level.

Aside obvious consequences on the consistency and genetic diversity of natural populations, when followed by release of individuals in the non-native range, pet trade can trigger several processes posing additional threats to wildlife: i) hybridization between native and translocated individuals [10,11]; ii) introduction of exotic parasites and pathogens [12]; iii) ecosystem imbalance [13,14]; iv) new biological invasions [15,16]. Therefore, limiting collection within the areas of origin, and correctly relocating confiscated individuals are activities of the utmost importance [17].

The Mediterranean land tortoises are known to be largely threatened by pet trade, especially in the Balkans [18–22], where the former Yugoslavia had an important role in tortoise exports during the past century [23–25]. According to the Federal Statistical Office, a total of 2,615 tons of tortoises were exported from the former Yugoslavia within a 41-year period during the 20th Century, approximating 2 million traded individuals [23]. The Hermann’s tortoise (*Testudo hermanni* Gmelin, 1789) has been particularly affected by this trade [26]. This species has its natural range spanning from Spain to the Balkans, mainly along the Mediterranean coastal regions, and in various Mediterranean islands. Two subspecies with clear genetic differences are commonly recognized (the eastern *T. h. boettgeri* and the western *T. h. hermanni*), and the geographic structure of the genetic variation in both subspecies, although with some under-sampled areas, has been assessed [27, 28]. Intensive harvesting for pet trade, especially before the 1980s when it was not banned yet [23], and releases of non-native individuals into local populations, are long-recognized threats for this species [26], along with habitat reduction [29]. As a consequence, *T. hermanni* is included in the list of the strictly protected species by the *Bern Convention on the Conservation of European Wildlife and Natural Habitat*, and the western subspecies *T. h. hermanni* is classified as “Endangered” by the Italian IUCN Red List of Vertebrates [30]. However, source and fate of illegally translocated individuals are still poorly assessed in vast portions of the species’ range.

In this paper, we test a small panel of microsatellite markers to investigate patterns of illegal translocations of *T. hermanni hermanni* among a large sample of individuals hosted in Italian seizure and recovery centers. To this end, we began by complementing previous assessments of population genetic structure of wild populations [28], with new data from previously unsampled geographic areas. Subsequently, we used information gathered from the Bayesian genetic clustering exercises to assign confiscated individuals to the most probable geographic area of origin.

## MATERIALS AND METHODS

### Sampling and laboratory methods

We collected 154 blood samples from wild *Testudo hermanni* individuals throughout mainland Italy, Sicily, Sardinia, Corsica and Lampedusa and 458 blood samples from confiscated tortoises kept in captivity by local authorities (e.g., the Carabinieri Corps) or animal conservation NGOs. Sampling sites of wild individuals and location of recovery centres are shown in Fig 1 and 2, respectively. Blood samples were taken from nape or coccygeal vein and about 75 μl were spotted on FTA^®^ Classic Cards (Whatman™, GE Healthcare) and stored at room temperature. Alternatively, whole blood (100μl – 1 ml) was treated with K3-EDTA and stored at -20° C. DNA was extracted from both FTA-Cards and whole blood using a solution of 5% Chelex^®^ 100 Resin (Bio-Rad, [31], see Supplementary Material). Initially, all individuals were genotyped at 9 microsatellite loci (Test10, Test56, Test71, Test76, Test88, Gal136, Gal75, Gal73, and Gal263) as in [28] (see also [32, 33]). However, two loci (Test88 and Gal73) yielded inconsistent reactions and were discarded from downstream analyses. Detailed protocols are provided as Supplementary Material. In order to combine our dataset with the dataset from Perez et al. [28] avoiding mislabelling of alleles, we re-genotyped selected samples from [28], and we recalibrated binning set and allele nomenclature to match their dataset. Fragment analysis of PCR products was performed by Macrogen Inc. on an ABI 3730xl Genetic Analyser (Applied Biosystems) with a 400HD size standard. Allele calling was performed with GENEMAPPER^®^ 4.1 checking electropherograms by eye. All electropherograms were scored by two persons and only concordant multilocus genotypes were retained for subsequent analyses.

**Figure 1.**
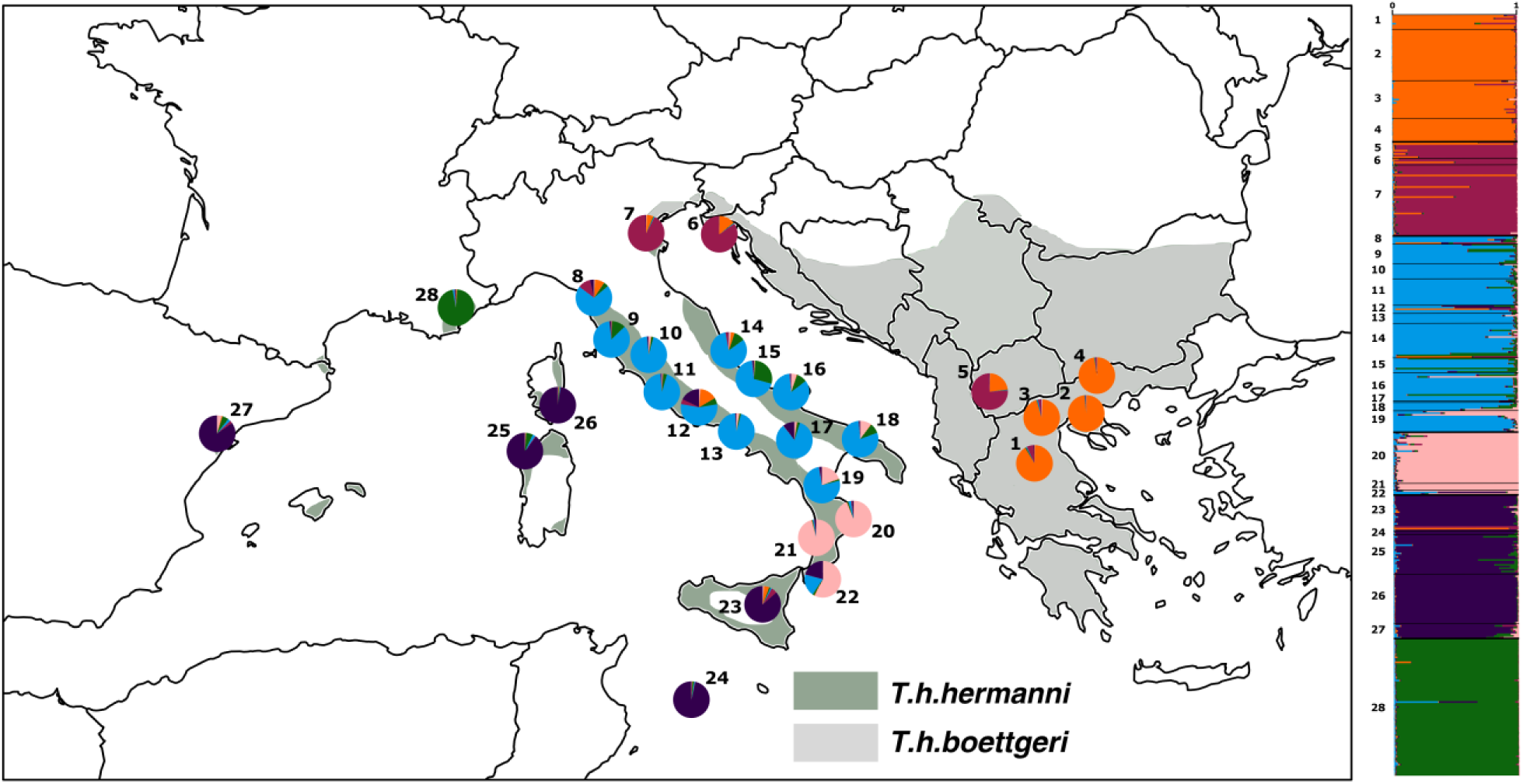
Genetic structure of wild *Testudo hermanni* populations estimated using STRUCTURE. The wild dataset resulted from the integration of samples collected in this study and individuals from [28]. Sampling locations and number of individuals sampled per sites (N) of the integrated dataset are: 1-Meteore [Greece] (N=9), 2-Epanomi [Greece] (N=31), 3-Aliki [Greece] (N=23), 4-Kerkini [Greece] (N=14), 5-Prespa Lake [Macedonia] (N=10), 6-Vodnjan [Croatia] (N=4), 7-Emilia-Romagna (N=43), *8-Tuscany North (N=5)*, *9-Tuscany South (N=12)*, *10-Lazio North (N=9)*, 11-Lazio Center (N=16), *12-Lazio South (N=5), 13-Campania North (N=6)*, *14-Abruzzo (N=21)*, *15-Molise (N=9)*, *16-Puglia North (N=17)*, *17-Campania Center (N=1)*, *18-Puglia South (N=5)*, *19-Calabria North (N=13)*, *20-Calabria Center-North (N=31)*, *21-Calabria Center-South (N=4)*, *22-Calabria South (N=3*), *23-Sicily (N=22)*, *24-Lampedusa (N=2)*, *25-Sardinia (N=24)*, *26-Corsica (N=30)*, 27-Ebro [Spain] (N=9), 28-Var [France] (N=83). In italic are shown new sampling sites from this study and locations whose sampling was increased from [28].

**Figure 2.**
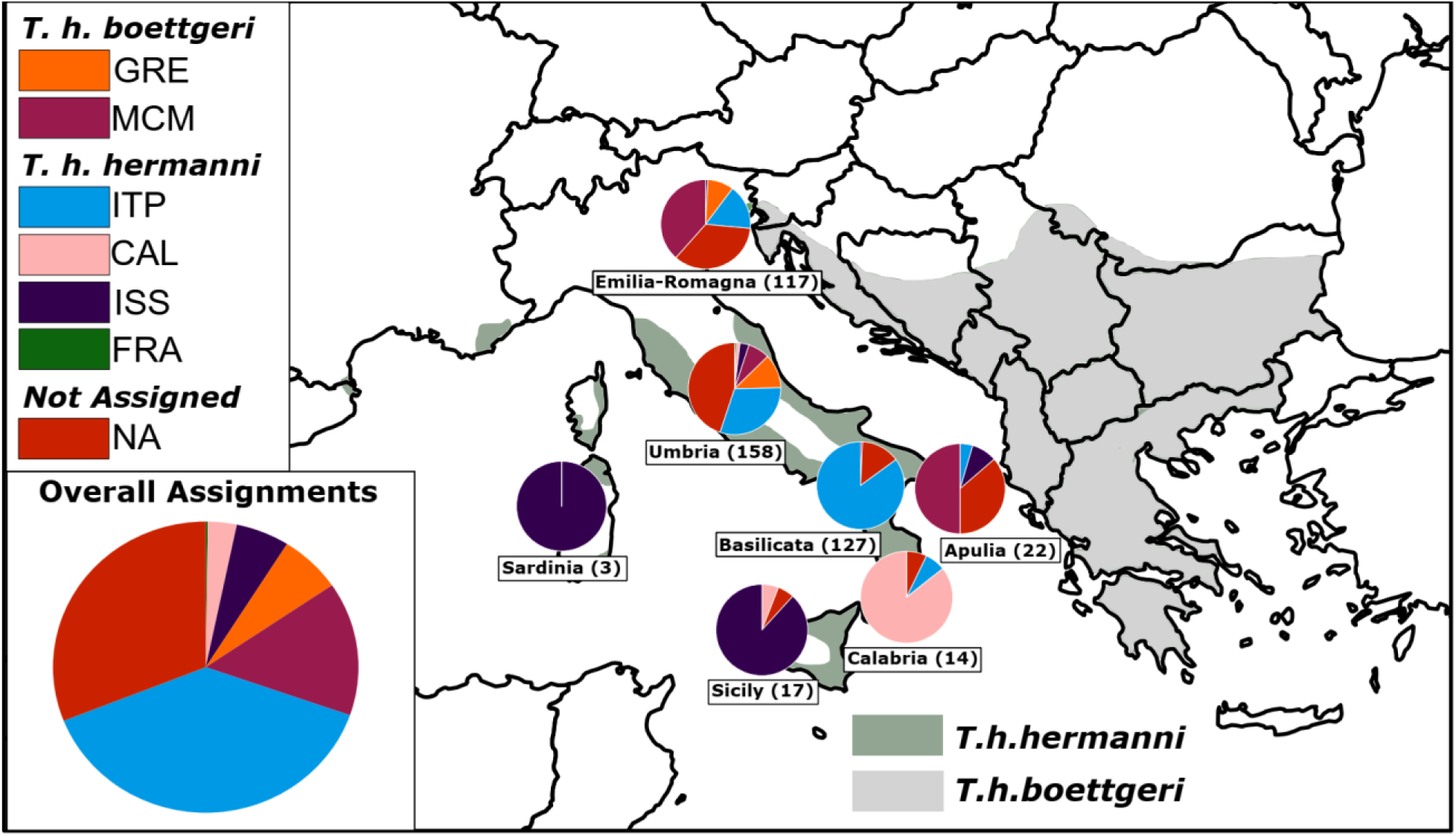
Geographic assignment of 458 confiscated samples from seven Italian seizure and recovery centres. Overall assignments are showed in the pie chart in the lower left corner. Local assignments for each recovery centre are showed in the pie charts on the map (in brackets the samples size). GRE = Greece; MCM = Bosco Mesola, Croatia and Macedonia; ITP = Italian Peninsula; CAL = Calabria; ISS = Mediterranean Islands and Spain; FRA = France; NA = not assigned samples.

### Genetic structure and reference database

As the first step to assess the area of origin of confiscated individuals, we carried out a population structure analysis of individuals that can be confidently considered as belonging to natural populations (hereon ‘wild’), in order to define possible source populations and to compile a reference database of individuals genuinely belonging to each identified population. The multilocus genotypes of the 154 newly collected wild tortoises were joined to the dataset from [28], excluding from the latter all the individuals that were reported to be migrant or 1^st^ and 2^nd^generation hybrids, and the admixed population of Bosco Nordio. The joint wild dataset consisted of 461 individuals (Fig 1). We performed the cluster analyses on the wild dataset using the Bayesian method implemented in STRUCTURE 2.3.4 [34]. Analyses were conducted choosing a model with admixture, uncorrelated allele frequencies, and a non-uniform ancestry prior ALPHA among clusters, as suggested by Wang [35] for uneven samplings. We run 20 replicates for each value of K from K=1 to K=12 (K is the number of inferred genetic groups), with 750000 MCMC after a burnin of 500000. Structure results were summarized and visualized with the web server CLUMPAK [36]. We used STRUCTURE HARVESTER [37] to infer the best value of K, based on both the probability of the data given K [34] and the Evanno approach [38].

The reference database of wild individuals was then prepared based on the following three-step analysis. First, we added to each wild individual the prior information about the genetic cluster most represented in the geographic area from which they were sampled. Second, we re-run STRUCTURE to identify migrants and hybrids, using the same parameters as above but fixing K at its optimal value (see results), and activating the USEPOPINFO option. Finally, all individuals that resulted as ‘non-pure’ in their respective geographic area (i.e. those individuals with less than 50% posterior probability to belong to their prior assigned cluster) were excluded from the reference database.

### Assignment of individuals of unknown origin

According to Manel and colleagues [39] fully Bayesian methods of assignment, as implemented in STRUCTURE, outperform partially Bayesian methods [40] with higher assignment rates and lower assignment error. However, this method considers all populations simultaneously with the drawback of assigning individuals to reference population even if the true population of origin is actually unsampled [39]. To overcome this problem, Manel and colleagues [39] suggests performing both fully Bayesian assignment tests and exclusion tests.

We performed assignment tests on 458 confiscated individuals with STRUCTURE using the POPFLAG for individuals in the reference database and activating the “update allele frequencies using only individuals with POPFLAG=1” option under a USEPOPINFO without admixture model. Other run parameters were the same as in the USEPOPINFO run described above. We assigned individuals to a source population when the probability of an individual to belong to that population was above 80%.

Exclusion tests were performed with the partially Bayesian exclusion method [41] implemented in GENECLASS2 [42]. We compared observed genotypes of confiscated individuals with an expected likelihood distribution of genotypes generated for each reference population by simulating 1000000 individuals with Monte Carlo resampling [43]. We excluded reference populations as the likely source of an individual when likelihood values were below 0.01.

## RESULTS

The analysis of the complete wild dataset indicated that K=2 and K=6 were the most supported numbers of clusters. The log probability of data increased sharply from K=1 to K=2 and then more slowly from K=3 to K=6 where it reached a plateau (see Supplementary Material Fig 1). The delta K analysis [38] provided two modes at K=2 and K=6, respectively. The first and most evident partition discriminated eastern and western subspecies (see Supplementary Material Fig 2), whereas the second mode at K=6 suggested a subdivision of *T. h. hermanni* in 4 groups and of *T. h. boettgeri* in 2 groups (Fig 1). The *T. h. hermanni* groups were Italian Peninsula (ITP) (all the populations from central and southern Italian Peninsula, except samples from central and southern Calabria), mainland France (FRA), Calabria (CAL) and Mediterranean islands (Sicily, Sardinia, Corsica, Pantelleria) joined with Spain (ISS). The *T. h. boettgeri* groups were Greece (GRE) and Bosco Mesola with Croatia and Macedonia (MCM). These results agree with the groups previously obtained by Perez et al. [27, 28, 44, 45], but with the additional CAL cluster, emerging from an area that was previously unsampled.

The analyses carried out in STRUCTURE using the prior population information allowed us to detect the presence of one hybrid and six migrant individuals among wild populations (Tab 1). While the hybrid was from an admixture area between two geographically contiguous clusters and one of the migrants was from the same subspecies, the other five migrants were from the other subspecies (four of them from spatially very distant clusters). Genotypes from these seven individuals were excluded from the reference database.

**Table 1.**
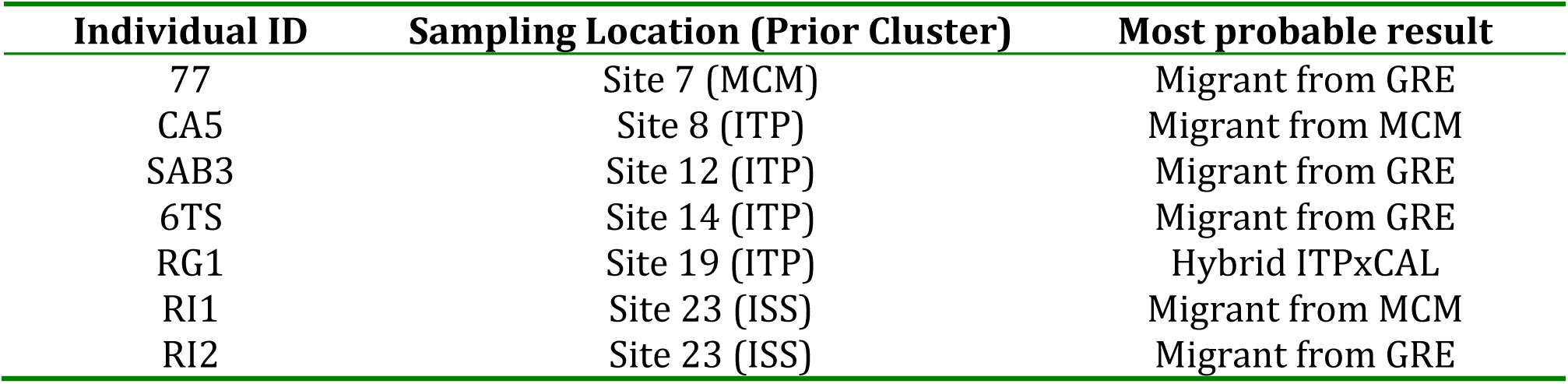
List of hybrid and migrant samples detected among wild populations. Sites are referred to the sampling locations (see fig. 1).

| Individual ID | Sampling Location (Prior Cluster) | Most probable result |
| --- | --- | --- |
| 77 | Site 7 (MCM) | Migrant from GRE |
| CA5 | Site 8 (ITP) | Migrant from MCM |
| SAB3 | Site 12 (ITP) | Migrant from GRE |
| 6TS | Site 14 (ITP) | Migrant from GRE |
| RG1 | Site 19 (ITP) | Hybrid ITPxCAL |
| RI1 | Site 23 (ISS) | Migrant from MCM |
| RI2 | Site 23 (ISS) | Migrant from GRE |

In order to assign the 458 confiscated individuals to the most probable geographic area of provenance, we used K=6 as the optimal K value, and q > 0.8 as the assignment threshold. Using these parameter values we were able to assign more than 90% of samples to one of the six clusters. When assigned individuals were downgraded to unassigned by the exclusion test (area of origin excluded with P<0.01), 38.7% of the confiscate tortoises were assigned to the ITP cluster, 14.8% to MCM, 6.5% to GRE, 5.7% to the ISS, 3.1% to CAL and 0.2% FRA, while 31% of the individuals were not assigned to any predefined cluster (NA). Decreasing the significance level of the exclusion test to 0.001 to avoid false positives in multiple testing decreased the fraction of unassigned individuals to 22%.

Most of the assigned tortoises belonged to the genetic cluster corresponding to the area where the captivity center was located (see Figure 2). However, we also found evidence of long distance translocations of individuals, especially in the centers along the Adriatic coast and facing the Balkan regions, known to be a source of illegal trades. In Apulia, for example, only one of the 14 assigned individuals belonged to the local ITP genetic cluster, whereas eleven of them were classified as MCM. In the Emilia-Romagna center, 60% of the assigned individuals were probably local or imported from Balkan areas genetically very similar (*Testudo hermanni boettgeri* MCM cluster), but about 25% and 14% and of them were classified as imported from Greece (GRE) or classified within the *Testudo hermanni hermanni* ITP cluster, respectively. In the Umbria centers, more than 20% of the assigned individuals had a Greek origin. On the other hand, when the small fraction of unassigned samples was excluded, more than 90% of the captive individuals from the Western and most Southern areas (Basilicata, Calabria, Sicily, and Sardinia) belonged to the local cluster.

## DISCUSSION

The main purpose of this work was to test a small panel of microsatellite markers potentially useful as a tool to identify the most probable geographic origin of *T. hermanni* tortoises, and to apply it to individuals of unknown origin confiscated because illegally owned or imported and currently hosted in Italian seizure and recovery centres. We found that this tool is able to assign a large fraction of individuals to specific macro-regions, thus contributing to forensic analysis and/or to projects of release in the wild of confiscated animals.

Results from the overall assignment tests showed that most of the assigned individuals were native of the Italian Peninsula (clusters ITP and CAL) or from clusters at least partially falling within national borders (clusters ISS and MCM). A significant 6.5% of genetically assigned tortoises hosted in Italian centres turned out to be of Greek origin, with evidence of long distance translocations. Only one individual was assigned to the French genetic cluster. These fractions, however, vary widely across seizure centres, with some of them hosting significant numbers of non-local individuals.

We found an overall 31% (22% using a more stringent criteria to exclude the source population identified by the assignment method) of captive individuals that were unassigned. This could be explained in the light of three main considerations. First, source populations of unassigned individuals may have remained unsampled. Our sampling scheme of the wild populations increased the coverage of the species range within the Italian borders (*T. h. hermanni*) [45] compared to previous studies [27, 28, 44, 46]. However, areas from outside this range remain poorly sampled, especially along the Balkan Peninsula, so it is possible that additional samples will improve the assignment performance in the future. Alternatively, an assignment approach combined with even more strict criteria to exclude populations may be used to assign additional individuals to populations which are genetically very similar, though distinct, from the source population. Second, unassigned individuals could be hybrids, due to mating occurred in captivity or in the wild (where introduced animals can be found) between individuals with different origin. In this case, assignment algorithms fail of course to assign individuals with high probability to a specific cluster. Third, different populations share relevant fractions of genetic variation, and therefore only more microsatellite markers could increase the discriminatory power of this assignment tool.

Our assessment of the genetic structure of wild populations confirmed the overall pattern found by Perez et al. [28], but also revealed further structure. Despite we used 7 microsatellite markers instead of the 9 used by Perez et al. [28], our results are fully consistent with theirs, showing 2 main genetic pools corresponding to the 2 recognized subspecies, and further structure within them. The increased sampling effort along the Italian Peninsula allowed us to recognize a further cluster in Calabria, a region recognized as glacial refugium and hotspot of genetic diversity for many temperate species [47–51]. The increased sampling effort on some Mediterranean islands (i.e., Sardinia and Lampedusa) confirmed the presence of a single insular genetic cluster.

The preliminary analysis performed on wild populations revealed the presence of six migrants and one hybrid among wild populations. While the hybrid individual found in the northern area of Calabria can reasonably be considered as a consequence of a natural admixture zone between Italian peninsular and Calabrian clusters, the presence of the migrants from far distant areas of origin could be explained by human-driven translocations. In particular, the presence of *T. h. boettgeri* individuals from Greece in wild populations along the Italian Peninsula and Sicily could be a consequence of the wide pet trade affecting this species, with hundreds of thousands tortoises collected mostly in south-eastern Europe between the 1960s and the 1980s and shipped to western Europe [18,52] or even of more ancient translocations [53]. This evidence clearly indicates that the escape or the release of non-endemic individuals among wild endemic populations is not so rare, with potential genetic and epidemiological implications.

A priority concern that motivated this study and requires urgent solutions is the management of the tortoises kept in captivity in seizure/recovery centres. These animals, usually confiscated from local authorities or found by private citizens far from natural populations and likely escaped from domestic contexts, cannot be released in nature without knowledge of their origin. Their number is increasing, with increasing problems related to their management and health condition. The assembly of a genetic reference database, and the assessment of the most probable geographic origin of captive tortoises, are fundamental steps towards the development of plans of reintroduction in the wild, which will not only reduce the problems and the costs associated with the captive animals, but also re-create wild populations in areas where this species was present in the past but is now extinct. In addition, the reference database represents a useful forensic tool to investigate the genotype of individuals when their declared origin is legally disputed.

Future efforts should be devoted to achieve higher geographic resolution of genetic population structure analyses, and to reduce the fraction of unassigned individuals. These goals could be achieved with one or both of the following strategies. First, to sample still poorly covered areas, in order to get a complete representation of the genetic variation in the whole species’ range. Second, to increase the number of informative genetic markers, possibly decreasing the costs. Next Generation Sequencing (NGS) technologies could help in this direction, allowing to develop a panel of diagnostic SNPs to be assessed with the increasingly cheap genotyping methods [1, 54].

## Supporting information

## ACKNOWLEDGMENTS

Animal handling and sample collection were allowed by the Ministero dell’Ambiente e della Tutela del Territorio e del Mare (0044068 - 4/12/2012-PNM-II; 0001805/PNM - 4/2/2015; ISPRA 68754/T-A31 – 28/11/2016) and the Regione Autonoma della Sardegna Prot.4749, Rep.N.73 07/03/2017. We would like to thank all the colleagues and friends who helped during fieldwork: Carabinieri per la Tutela dell’Ambiente; Centro Regionale di Recupero degli animali selvatici di Bonassai, Centro Recupero Fauna Selvatica Bosco di Ficuzza; Centro Recupero Animali Selvatici Formichella; Centro Recupero Fauna Selvatica “Stretto di Messina”; Centro Recupero Animali Selvatici Provinciale di Policoro; Parco Nazionale dell’Asinara - Area Marina Protetta “Isola dell’Asinara”; Parco Nazionale dell’Aspromonte, Parco Nazionale del Circeo, Riserva Naturale Regionale Lecceta di Torino di Sangro; Riserva Naturale Bosco della Mesola - Parco Delta del Po; Oasi WWF “Lago di Conza. Finally, a special thank to Federica Baldo and Giulia Fabbri for their help and assistance, and Giovanni Nobili and all the Carabinieri Forestali at Punta Marina (Ravenna) and Bosco Mesola (Ferrara) for the continuous support and help. Funding was also provided by the University of Ferrara and the Ufficio Territoriale Carabinieri per la Biodiversità (Punta Marina).

