## Supplementary material for "Who’s who in the western Hermann’s tortoise conservation: a STR toolkit and reference database for wildlife forensic genetic analyses"

### MATERIALS AND METHODS

#### DNA extraction

**FTA-Cards**. One spot of blood was punched from FTA Classic Crads with a Harris Uni-Core puncher (1.2 mm), placed in a 1.5 ml microcentrifuge tube and macerated with a needle. The disc sample was washed twice with 1 ml of sterile deionized water, incubating for 10 min with occasional vortexing. Samples were then centrifuged for 3 min at 20000 x g, the supernatant was removed and discarded. Next, 200 µl of 5% Chelex 100 suspension was added to the sample and incubated at 56° C for 20 min. Samples were then vortexed for 15 sec and incubated at 100° C for 8 min. After incubation samples were vortexed again for 15 sec and centrifuged for 3 min at 20000 x g. The supernatant was transferred to a new tube and stored at -20° C.

**Whole blood**. 3 µl of blood were added to 1 ml of sterile deionized water in a 1.5 ml microcentrifuge tube, mixed and incubated at room temperature for 15-30 min to wash. Samples were then centrifuged for 3 min at 10000 x g. Supernatant were removed and discarded, leaving about 20-30 µl of sample in the tube. Next, 5% of Chelex suspension was added to the samples until to 200 µl of final volume and incubated at 56° C for 20 min. Samples were then vortexed for 5 sec and incubated at 100° C for 8 min. After incubation samples were vortexed again for 5 sec and centrifuged for 3 min at 10000 x g. The supernatant was transferred to a new tube and stored at -20° C.

#### Microsatellite PCR

**Test10**: forward primer was labelled with 6-FAM; reaction mix (10 µl) contained 2 µl of DNA, 0.5 µM of each primer, 2.5 mM $MgCl_2$, 0.2 mM of each dNTP, GoTaq Flexi Buffer 1X and 0.6 U of GoTaq G2 Hot-Start DNA Polymerase (Promega); PCR cycling included an initial denaturation at

94° C for 2 min, 35 cycles of 20 sec at 94° C, 10 sec at 53° C, 20 sec at 65° C and a final extension at 65° C for 5 min.

**Test71**: forward primer was labelled with 6-FAM; reaction mix (10 µl) contained 2 µl of DNA, 0.6 µM of each primer, 2.5 mM $MgCl_2$, 0.2 mM of each dNTP, GoTaq Flexi Buffer 1X and 0.75 U of GoTaq G2 Hot-Start DNA Polymerase (Promega); PCR cycling included an initial denaturation at 94° C for 2 min, 37 cycles of 30 sec at 94° C, 1 min at 56° C, 1 min at 72° C and a final extension at 72° C for 5 min.

**Gal75**: forward primer was labelled with 6-FAM; reaction mix (10 µl) contained 2 µl of DNA, 0.24 µM of each primer, 3 mM $MgCl_2$, 0.2 mM of each dNTP, GoTaq Flexi Buffer 1X and 0.6 U of GoTaq G2 Hot-Start DNA Polymerase (Promega); PCR cycling included an initial denaturation at 94° C for 5 min, 15 cycles of 30 sec at 94° C, 30 sec at 60° C (decreasing the annealing temperature of 1° C at each cycle), 30 sec at 72° C, 20 cycles of 30 sec at 94° C, 30 sec at 45° C, 30 sec at 72° C, and a final extension at 72° C for 7 min.

**Gal263**: forward primer was labelled with HEX; reaction mix (10 µl) contained 2 µl of DNA, 0.5 µM of each primer, 1.5 mM $MgCl_2$, 0.2 mM of each dNTP, GoTaq Flexi Buffer 1X and 0.5 U of GoTaq G2 Hot-Start DNA Polymerase (Promega); PCR cycling included an initial denaturation at 94° C for 3 min, 35 cycles of 30 sec at 94° C, 30 sec at 50° C, 30 sec at 72° C and a final extension at 72° C for 3 min.

**Test56**: forward primer was labelled with HEX; reaction mix (10 µl) contained 2 µl of DNA, 0.5 µM of each primer, 2.5 mM $MgCl_2$, 0.2 mM of each dNTP, GoTaq Flexi Buffer 1X and 0.6 U of GoTaq G2 Hot-Start DNA Polymerase (Promega); PCR cycling included an initial denaturation at 94° C for

2 min, 35 cycles of 20 sec at 94° C, 10 sec at 56° C, 20 sec at 65° C and a final extension at 65° C for 5 min.

**Test76**: forward primer was labelled with TAMRA; reaction mix (10 µl) contained 2 µl of DNA, 0.5 µM of each primer, 2.5 mM $MgCl_2$, 0.2 mM of each dNTP, GoTaq Flexi Buffer 1X and 0.6 U of GoTaq G2 Hot-Start DNA Polymerase (Promega); PCR cycling included an initial denaturation at 94° C for 2 min, 35 cycles of 20 sec at 94° C, 10 sec at 58° C, 20 sec at 65° C and a final extension at 65° C for 5 min.

**Gal136**: forward primer was labelled with 6-FAM; reaction mix (10 µl) contained 2 µl of DNA, 0.24 µM of each primer, 3 mM $MgCl_2$, 0.2 mM of each dNTP, GoTaq Flexi Buffer 1X and 0.6 U of GoTaq G2 Hot-Start DNA Polymerase (Promega); PCR cycling included an initial denaturation at 94° C for 5 min, 15 cycles of 30 sec at 94° C, 30 sec at 60° C (decreasing the annealing temperature of 1° C at each cycle), 30 sec at 72° C, 20 cycles of 30 sec at 94° C, 30 sec at 45° C, 30 sec at 72° C, and a final extension at 72° C for 7 min.

PCR products were assembled in multiplex post-PCR: one including Test76, Gal263, Gal75, Test71, Test10 and another including Gal136 and Test56.

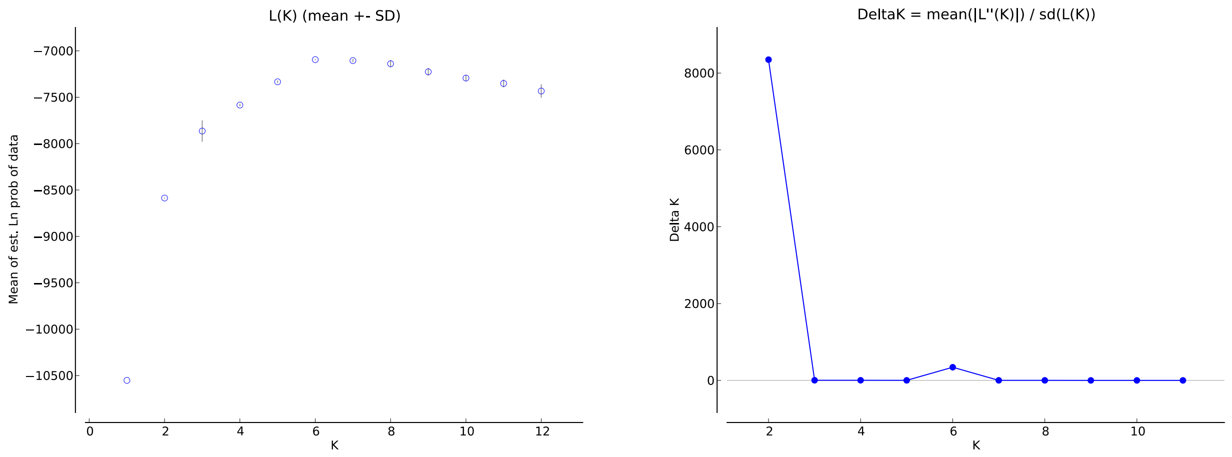


**Figure S1.** Probability of number of clusters (K) for *Testudo hermanni* sampled at 28 localities, analysed at seven microsatellite loci, using STRUCTURE HARVESTER [1]. The Ln likelihood value (figure to the left) as described by [2] and the Delta K method (figure to the right) by [3].
